# Sublethal effects of the neonicotinoid insecticide thiamethoxam on the transcriptome of the honeybee (*Apis mellifera*)

**DOI:** 10.1101/114256

**Authors:** Teng-Fei Shi, Yu-Fei Wang, Lei Qi, Fang Liu, Lin-Sheng Yu

**Affiliations:** School of Plant Protection, Anhui Agricultural University, Hefei 230036, China; School of Animal Science and Technology, Anhui Agricultural University, Hefei 230036, China

**Keywords:** Thiamethoxam, honeybees (*Apis mellifera*), RNA-Seq, differential gene expression

## Abstract

Neonicotinoid insecticides are now the most widely used insecticides in the world. Previous studies have indicated that sublethal doses of neonicotinoids impair learning, memory capacity, foraging and immunocompetence in honeybees (*Apis mellifera*). Despite this, few studies have been carried out on the molecular effects of neonicotinoids. In this study, we focus on the second-generation neonicotinoid thiamethoxam, which is currently widely used in agriculture to protect crops. Using high-throughput RNA-Seq, we investigated the transcriptome profile of honeybees after subchronic exposure to thiamethoxam (10 ppb) over 10 days. In total, 609 differentially-expressed genes (DEGs) were identified, of which 225 were up-regulated and 384 were down-regulated. The functions of some DEGs were identified, and GO enrichment analysis showed that the enriched DEGs were mainly linked to metabolism, biosynthesis and translation. KEGG pathway analysis showed that thiamethoxam affected biological processes including ribosomes, the oxidative phosphorylation pathway, tyrosine metabolism pathway, pentose and glucuronate interconversions and drug metabolism. Overall, our results provide a basis for understanding the molecular mechanisms of the complex interactions between neonicotinoid insecticides and honeybees.

**Summary statement:** *NR1, Cyp6as5, nAChRa9* and *nAChRβ2* were up-regulated in honeybees exposed to thiamethoxam, while *CSP3, Obp21, defensin-1, Mrjp1, Mrjp3* and *Mrjp4* were down-regulated.

## 1. Introduction

Honeybees (*Apis mellifera,* L.) have a high social and economic value since they produce various substances such as honey and also play an important role in pollination and agricultural production (Breeze *et al.*, 2011). In recent years, attention has been paid to the large decrease to global apiculture (Neumann and Carreck, 2010; Potts et al., 2010; Van Engelsdorp *et al.*, 2010; Chauzat *et al.*, 2013) but the reasons are still poorly understood. Recent studies have however suggested that the decrease could be due to the widespread use of insecticides (Johnson *et al.*, 2010; Goulson *et al.*, 2015; Schmuck and Lewis, 2016).

Recently, there have been far-reaching changes in the insecticide market. Many of the traditional insecticides, *e.g.* organophosphorus and pyrethroids, have been replaced by systemic insecticides, especially neonicotinoids. Neonicotinoids act on the insect nervous system mainly through agonistic action on nicotinic acetylcholine receptors (nAChRs) (Brown *et al.*, 2006), and since they have low mammalian toxicity (Tomizawa and Casida, 2005) they are widely used for controlling insect pests. Neonicotinoids are commonly applied as seed coatings or as foliar sprays on crops. Once absorbed into the plant, neonicotinoids can translocate to dew drops, nectar and pollen of crops during florescence (Krupke *et al.*, 2012; Stoner and Eitzer, 2012). The contaminated nectar and pollen may be consumed by foragers (Goulson, 2013) or taken to the nest for long-term storage where they are eaten by the young adults and larvae (DeGrandi-Hoffman *et al.*, 2000; Cresswell, 2011). Recent studies have detected various neonicotinoids in bee products, *e.g.* honey, pollen and beeswax (Stoner and Eitzer, 2012; Codling *et al.*, 2016; Sánchez-Hernández *et al.*, 2016), meaning that the neonicotinoids can have chronic effects.

Even though several neonicotinoids, including thiamethoxam, imidacloprid and clothianidin, have been found to be highly toxic to honeybees (Laurino *et al.*, 2011), they are not acutely lethal at field levels (Blacquière *et al.*, 2012). Nevertheless, there are considerable chronic and sublethal effects, including impairment to the brain, mushroom body and midgut (Catae *et al.*, 2014; Oliveira *et al.*, 2014; Peng and Yang, 2016) and decreased learning and memory capacity (Aliouane *et al.*, 2009; Mengoni and Farina, 2015; Alkassab and Kirchner, 2016). Evidence from semi-field or field research indicated that neonicotinoids negatively affect foraging activity and homing flight (Henry *et al.*, 2012; Fischer *et al.*, 2014; Tison *et al.*, 2016). Moreover, neonicotinoids have been found to affect honeybee immunocompetence (Brandt *et al.*, 2016) and increase the risk of other stressors such as pathogens (Pettis *et al.*, 2013; Alburaki *et al.*, 2015).

Despite the implications for honeybee colonies, little research has so far been carried out into the molecular effects of neonicotinoids. Christen *et al.* (2016) found that exposure to neonicotinoids changed the transcription of *AChR*α*1* and *2, creb, pka* and *vitellogenin* in the brain of honeybees. The latest research from this group (Christen *et al.*, 2017) showed that binary mixtures of neonicotinoids lead to different transcriptional changes in *nAChR* subunits and *vitellogenin* than single neonicotinoids, and that transcription was most strongly induced by thiamethoxam.

In the current study, we focused on the second-generation neonicotinoid thiamethoxam (Maienfisch *et al.*, 2003). Using high-throughput RNA-Seq, we investigated the transcriptome profile of honeybees after exposure to a sublethal concentration (10 ppb) for 10 days. The transcriptome profiles were then systematically analyzed by differential gene expression, Gene Ontology (GO) categories and Kyoto Encyclopedia of Genes and Genomes (KEGG) pathways. Our study aims to provide a basis to explore the molecular mechanisms of thiamethoxam and contribute to the understanding of the decline in honeybee populations.

## 2. Results

### 2.1. Raw read processing and quantitative gene expression

Using high-throughput RNA-Seq, six libraries (SCTH_1-3 and CK_1-3) were created from the two treatments (SCTH and CK). In total, SCTH_1, SCTH_2, SCTH_3, CK_1, CK_2 and CK_3 generated 43,672,706, 42,805,654, 43,630,710, 43,594,286, 44,650,260 and 43,379,868 usable reads, respectively. After mapping to the reference genome (NCBI: Amel_4.5) and the junction database, 38,527,526, 38,026,904, 38,277,904, 39,258,011, 40,153,044 and 37,645,388 total mapped reads were acquired, and the numbers of uniquely mapped reads were 37,673,745, 37,163,425, 37,440,667, 38,121,707, 38,960,068 and 36,686,103. Among these unique reads, 77.58-81.93% were mapped to exon regions (Table S1).

We calculated gene activity by counting the reads that mapped to exon regions (≥ 3 per gene). The average number of genes expressed in SCTH and CK libraries was 11,426 and 11,330 respectively; and 11,150 genes were expressed in both groups (Fig. 1). We also divided gene expression levels into five grades according to their RPKM (Reads Per Kilo bases per Million reads) values (Table S2). In each library, 30.22–31.79% of the reads had RPKM values < 1, 11.49–12.56% had RPKM values of 1-3, 28.20–29.17% had RPKM values of 3-15, 18.36–20.86% had values of 15-60 and 8.23–8.95% had RPKM values > 60. These results showed that a small number of genes were expressed at very high levels but the majority were expressed at low levels, indicating that the distribution of our DGE dataset was normal.

**Figure 1.**
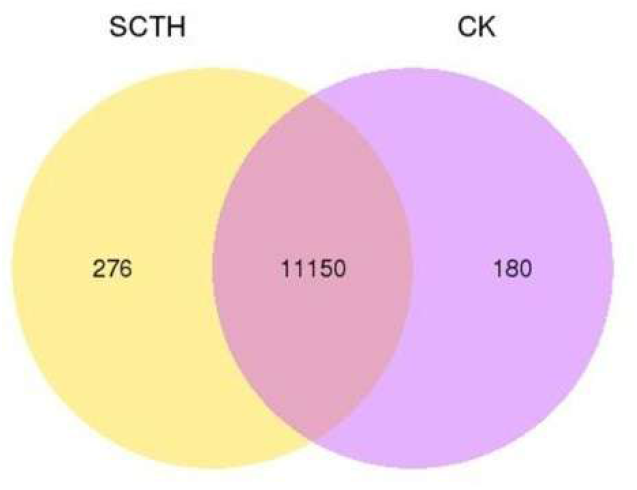
The average number of genes specifically expressed in SCTH and CK libraries, shown as number of genes expressed in each class. Honeybees exposure to sublethal concentration of thiamethoxam (10ppb) over 10 days, with acetone as the control check. Each library (SCTH or CK) involved three biological replicates of 15 bees each. SCTH: sublethal concentration of thiamethoxam; CK: control check.

### 2.2. Differentially-expressed genes between SCTH and CK honeybees

In total, 609 differentially-expressed genes (DEGs) were detected, of which 225 (45.2%) were up-regulated and 334 (54.8%) were down-regulated in the SCTH bees compared with CK bees (Fig. 2, Table S3). A list of the 20 genes with the most significant differential expression is shown in Table S4 and of these, 17 were down-regulated and 3 were up-regulated in SCTH honeybees. The 67 confirmed DEGs are listed in Table S5, and the others have been designated as hypothetical proteins. We focused here only on those that had previously been confirmed (Table 1).

**Table 1.**
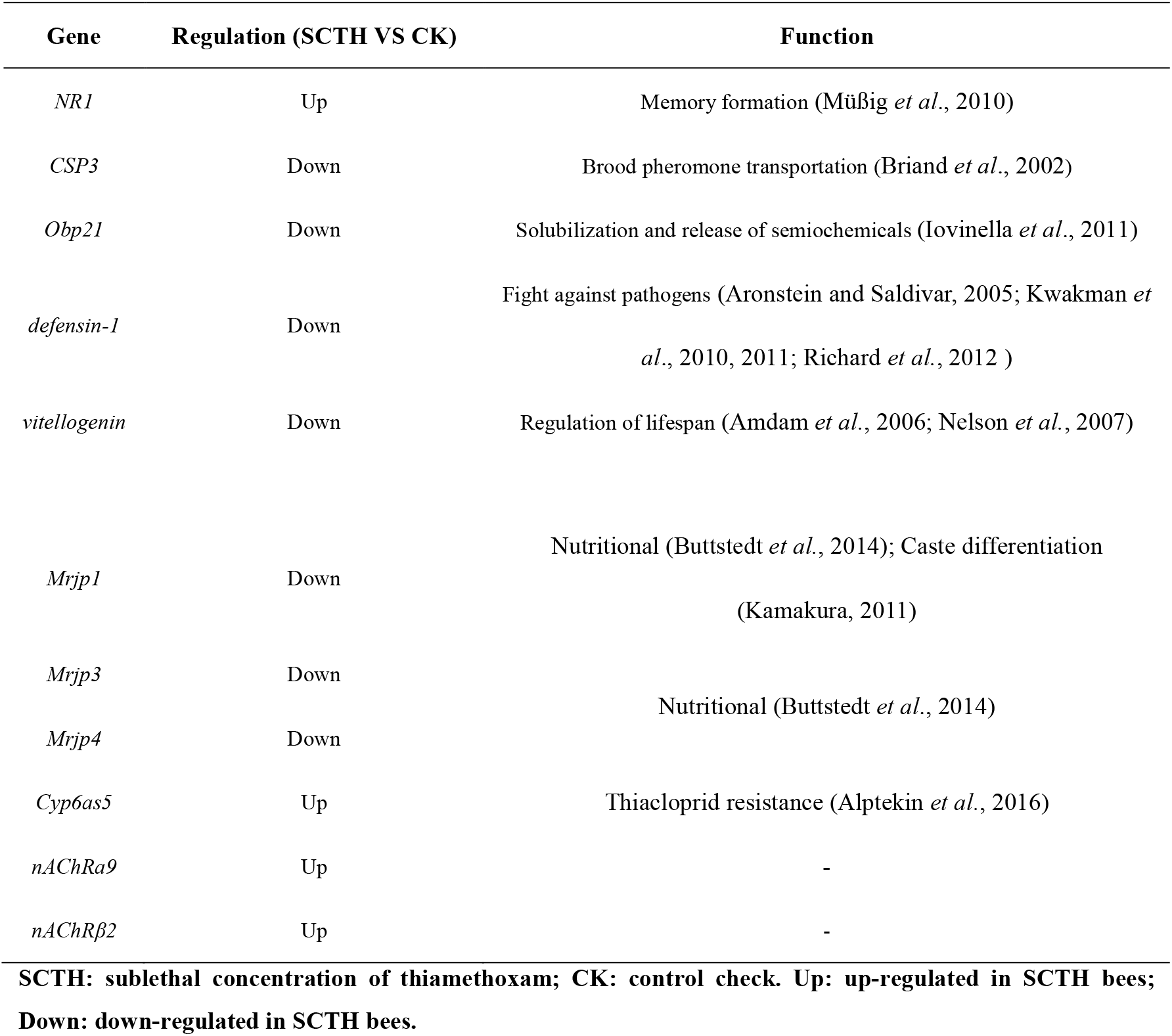
Information on selected differentially-expressed genes between SCTH and CK honeybees (*Apis mellifera*), corrected P-value ≤ 0.05.

We found that 10 ribosomal protein (RP) genes (*RpL37, RpS8, RpSA, RpL32, RpL18A, RpS3A, RpS6, RpS12, RpL13* and *RpL19*) have high expression levels, but were down-regulated in SCTH honeybees. In contrast, two nicotinic acetylcholine receptors (nAChRs) subunits, *nAChR*α*9* and *nAChR*β*2,* were up-regulated along with cytochrome P450 6AS5 (*Cyp6as5*).

Some genes, for example *defensin 1, vitellogenin,* LOC725387 and LOC406093, all have very high expression levels in both SCTH and CK honeybees, with intensity read copy counts of > 10,000. *Defensin1, vitellogenin* and LOC406093 were down-regulated in SCTH honeybees, whereas LOC725387 was up-regulated.

Odorant-binding proteins (OBPs) and chemosensory proteins (CSPs) are believed to be involved in odor recognition and chemical communication (Pelosi *et al,* 2006; Sanchez-Gracia *et al,* 2009). The genes *Obp3, Obp17, Obp21* and *CSP3* all showed significantly decreased expression in SCTH honeybees. Moreover, three major royal jelly protein (MRJP) coding genes, *Mrjp, Mrjp2* and *Mrjp3,* were down-regulated in the SCTH group. Although the expression level of a memory-related gene, *NMDA receptor 1* (*NR1*), was relatively low, it was differentially expressed between SCTH and CK honeybees.

**Figure 2.**
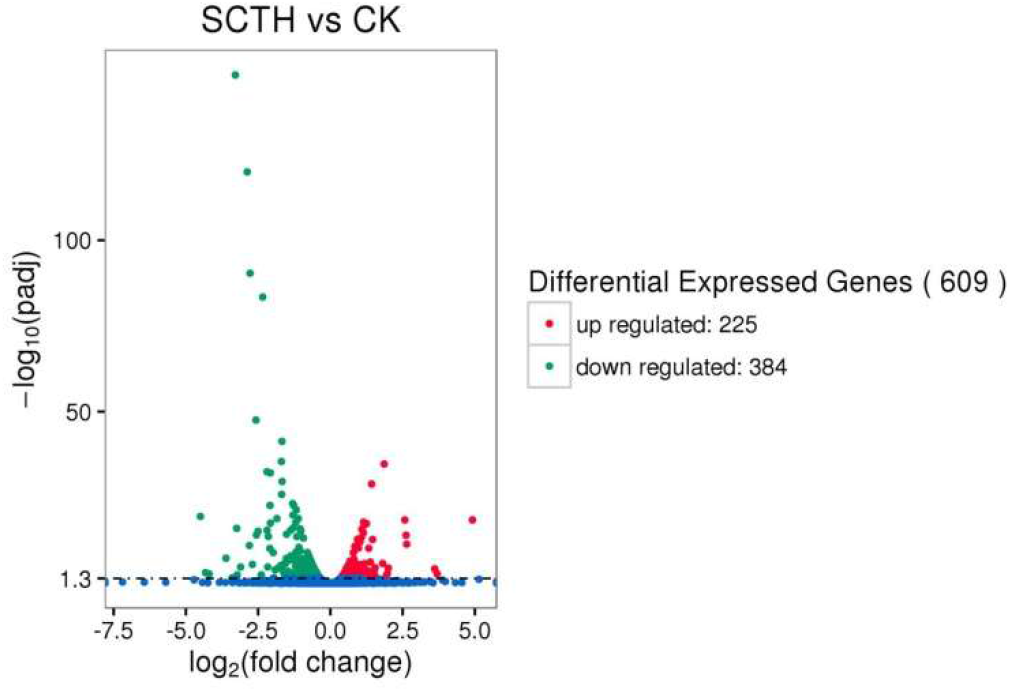
Volcano plot of differentially-expressed genes between SCTH and CK honeybees (*Apis mellifera*). Each treatment (SCTH or CK) involved three biological replicates of 15 bees each. Genes with an adjusted P-value (padj) ≤ 0.05 (FDR correction method) were considered differentially expressed between SCTH and CK bees. Red points: up-regulated genes in SCTH bees; Green points: down-regulated genes in SCTH bees; Blue points: no significant difference. SCTH: sublethal concentration of thiamethoxam; CK: control check.

### 2.3. Gene ontology (GO) enrichment analysis

A total of 445 DEGs were enriched for GO terms, including 167 up-regulated and 278 down-regulated genes in SCTH bees (Table S6), and the top 30 most enriched terms are shown in Fig 3. The genes were divided into three classes: molecular function, cellular components and biological process. Based on the GO terms for biological process, we found that most genes were enriched for translation, various metabolic and biosynthetic processes, such as protein metabolism, cellular protein metabolism, single-organism metabolism, cellular biosynthetic, cellular macromolecule biosynthetic, macromolecule biosynthetic and organic substance biosynthetic (Fig 3). The main DEGs that were enriched coded for cellular components, including the ribosome, ribonucleoprotein complex, ribosomal subunit and ribosomal subunit. Most of the genes were down-regulated in SCTH honeybees (Fig 3, Table S6). In terms of molecular function, the DEGs played roles in structural constitutent of ribosome, structural molecule activity and oxidoreductase activity (Fig 3).

**Figure 3.**
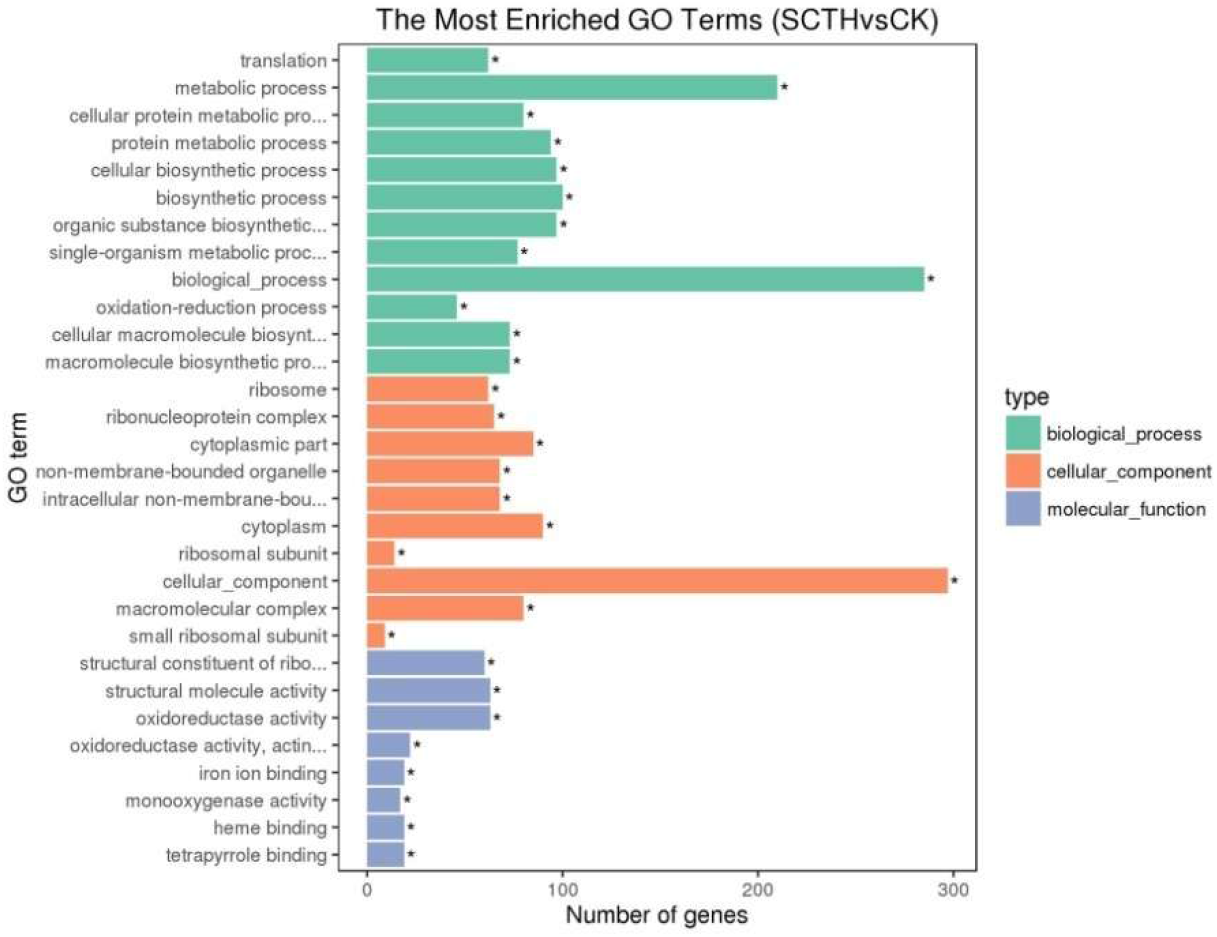
GO enrichment analysis of the differentially-expressed genes (DEGs) between SCTH and CK honeybees (*Apis mellifera*). Green bars: DEGs were enriched for biological process; Orange bars: DEGs were enriched for cellular component; Purple bars: DEGs were enriched for molecular function. Asterisk indicates GO terms were significantly enriched by DEGs (corrected *P*-values ≤ 0.05, FDR correction method). SCTH: sublethal concentration of thiamethoxam; CK: control check.

### 2.4 KEGG pathway analysis

The KEGG database (http://www.genome.jp/kegg) was used to assign functional annotations to the DEGs. A total of 377 DEGs were identified and mapped to 75 pathways in the KEGG pathway database (Table S7), including 104 up-regulated and 273 down-regulated genes. Among these pathways, five were significantly enriched with a corrected *P*-value ≤ 0.05 (Table 2). These included the regulation of most genes related to ribosomes, oxidative phosphorylation, tyrosine metabolism, pentose and glucuronate interconversion and drug metabolism.

**Table 2.** The five significantly enriched pathways, corrected *P*-value ≤ 0.05.

| Pathways | Pathway ID | Gene number | Corrected $P$-value |
| --- | --- | --- | --- |
| Ribosome | ame03010 | 64 | 1.1001522981e-25 |
| Oxidative phosphorylation | ame00190 | 19 | 0.0286819305139 |
| Tyrosine metabolism | ame00350 | 6 | 0.042425360978 |
| Pentose and glucuronate interconversions | ame00040 | 7 | 0.0286819305139 |
| Drug metabolism - other enzymes | ame00983 | 7 | 0.0369487673874 |

## 3. Discussion

The neonicotinoid insecticide thiamethoxam is highly toxic to honeybees with LD_50_ values in the range of a few ng per bee. The sublethal effects of thiamethoxam on honeybees have been extensively studied at many different physiological levels (Aliouane *et al.*, 2009; Henry *et al.*, 2012; Catae *et al.*, 2014; Oliveira *et al.*, 2014; Alburaki *et al.*, 2015). Here we identified 609 differentially-expressed genes (DEGs) in honeybees on exposure to sublethal concentration of thiamethoxam, including 225 up-regulated genes and 384 down-regulated genes.

The results of GO and KEGG analysis showed that the regulation of many DEGs related to metabolism, biosynthesis and translation, the ribosome pathway, oxidative phosphorylation pathway, tyrosine metabolism pathway and pentose and glucuronate interconversions. This suggests that thiamethoxam mainly affects physiological and developmental processes in honeybees.

Nicotinic acetylcholine receptors (nAChRs) are targets of neonicotinoids and their induction can produce overt effects such as decreased memory and locomotor capacity (Aliouane *et al.*, 2009; Charreton *et al.*, 2015). Our study showed that subchronic exposure to thiamethoxam increases the expression of two nAChRs subunits, *nAChRα9* and *nAChRβ2.* These results are consistent with previous research that thiamethoxam up-regulates expression of *nAChRα1* and *nAChRα2* (Christen *et al.*, 2016; Christen *et al.*, 2017). This indicates a compensatory reaction to the functional loss of nAChRs due to neonicotinoids (Christen *et al.*, 2016).

NMDA glutamate receptors (NMDARs) are composed of NR1 and NR2 subunits and are known to play an important role in memory formation (Kandel, 2001): injection of NMDA receptor antagonists led to an impairment of long-term memory (Si *et al.*, 2004). The NR1 subunit is expressed throughout the honeybee brain and plays a critical role in the functional expression of NMDARs (Zannat *et al.*, 2006). Müβig *et al.* (2010) reported that inhibiting expression of the *NR1* subunit in the honeybee brain can impair the formation of mid-term and early long-term memory. Thus, the thiamethoxam-induced alteration of *NR1* in our study might partly explain the effects on memory formation.

There are two known classes of small soluble proteins in the chemosensilla of insects: odorant-binding proteins (OBPs) and chemosensory proteins (CSPs). These are believed to be involved in odor recognition and chemical communication (Pelosi *et al.*, 2006; Sanchez-Gracia *et al.*, 2009), and CSP3 is thought to play a role in brood pheromone transportation (Briand *et al.*, 2002). OBP21 can bind the main components of queen mandibular pheromone as well as farnesol, a compound used as a trail pheromone (Iovinella *et al.*, 2011). The down-regulation of *CSP3* and *Obp21* seen in our study suggested a reduced chemosensory ability in challenged honeybees. This is generally consistent with recent research which indicated that a sublethal dose of the neonicotinoid imidacloprid decreased the binding affinity of OBP2 to a floral volatile, β-ionone, in Asiatic honeybees (Li *et al.*, 2015).

Neonicotinoids could also affect the immunocompetence of honeybees. A recent study reported that exposure to field-realistic concentrations of imidacloprid decreased hemocyte density, the encapsulation response and antimicrobial action (Brandt *et al.*, 2016). Antimicrobial peptides are a key component of honeybee innate immunity and include defensin, which is coded by two different defensin genes (*defensin-1* and *defensin-2*) in the honeybee genome (Evans *et al.*, 2006). *Defensin-1* has been found to be up-regulated following bacterial challenge (Aronstein and Saldivar, 2005; Richard *et al.*, 2012), suggesting that it is important in pathogen defense. Kwakman *et al.* (2010, 2011) found that *defensin-1* is a key antimicrobial compound of honey. The thiamethoxam-induced down-regulation of *defensin-1* found in our experiment may be involved in reducing the effectiveness of the immune system.

Vitellogenin is also a component of the defense mechanism since it acts as a free-radical scavenger to protect honeybees from oxidative stress (Amdam *et al.*, 2006; Seehuus *et al.*, 2006). Down-regulation of vitellogenin can shorten the lifespan of honeybees (Nelson *et al.*, 2007), so the thiamethoxam-induced decrease in *vitellogenin* expression in our experiment might contribute to reducing honeybee longevity.

Nutrition also plays an important role in honeybee development and longevity. Royal jelly (RJ) is a natural source of nutrients such as essential amino acids, lipids, vitamins and acetylcholine. Major Royal Jelly Proteins (MRJPs) constitute around 90% of total RJ protein. So far, nine MRJPs (MRJP1-MRJP9) and their encoding genes (*Mrjp1-Mrjp9*) have been identified, and MRJPs1-MRJPs5 have been suggested to have a nutritional function due to their abundance in RJ protein (Buttstedt *et al.*, 2014). MRJP1 is the predominant MRJP and is more highly expressed in the hypopharyngeal glands than in other body parts. An MRJP1 monomer, royalactin, was found to drive queen development through an Egfr-mediated signaling pathway (Kamakura, 2011). The honeybees used in our experiments were all young worker bees which can produce RJ in the hypopharyngeal glands. The thiamethoxam-induced down-regulation of *mrjp1, mrjp3* and *mrjp4* might decrease MRJP synthesis and indirectly cause a nutrition reduction in the lavae and queen.

However, just like other insects, honeybees have systems to detoxify insecticides. Cytochrome P450 monooxygenases (P450s) are the main detoxification enzymes which play an important role in the detoxification and metabolism of xenobiotics and insecticides. A P450 gene, *CYP6as5,* was induced in our experiment. This gene is a member of the CYP6AS subfamily, which have been implicated in the metabolism of xenobiotics in honey and pollen (Mao *et al.*, 2009; Johnson *et al.*, 2012). A recent study showed that *CYP6as5* was also significantly over-expressed in thiacloprid-treated bees compared with untreated controls, and induced thiacloprid insensitivity (Alptekin *et al.*, 2016). The P450 gene, *CYP6as5,* seems to play a central role in neonicotinoid insecticide resistance in honeybees.

In summary, using high-throughput RNA-Seq and analysis of differential gene expression, we detected 609 differentially-expressed genes in honeybees (*Apis mellifera*) after challenge with a sublethal concentration of thiamethoxam. We identified several genes involved in various physiological functions, but further studies are needed to confirm the results of this analysis. GO terms and KEGG pathways were also used to further understand the function of these genes. Our results provide a reference for understanding the molecular mechanisms of the complex interactions between neonicotinoid pesticides and honeybees.

## 4. Materials and Methods

### 4.1. Honeybee rearing

Two frames with sealed broods near adult emergence were taken from a healthy colony at the Institute of Apiculture Research of Anhui Agriculture University (Hefei, China). The population had not previously been exposed to pesticide. From July to September 2016 the frames were held in an incubator under the following conditions: 35 ± 1°C, a relative humidity (RH) of 50 ± 10% and in darkness. We obtained the newly-emerged honeybees and put them into cages (11 × 10 × 8 cm). They were fed with bee bread collected from the same apiary, sucrose-water solution (1:1 w:v), and maintained for three days at 30 ± 1 °C, a RH of 70 ± 10% and in darkness.

### 4.2. Thiamethoxam preparation and exposure

The residues of thiamethoxam in trapped pollen generally ranges from 0.6 to 53.3 ppb (Mullin *et al.*, 2010; Krupke *et al.*, 2012; Stoner and Eitzer, 2012; Codling *et al.*, 2016; Sánchez-Hernández *et al.*, 2016), and in honey from 2.5 to 17.2 ppb (Codling *et al.*, 2016; Sánchez-Hernández *et al.*, 2016). Based on this, a field-realistic level of 10 ppb (Stanley and Raine, 2016) was selected as the sublethal concentration. A stock solution of thiamethoxam (> 99% purity, 1000 ppm) was obtained from J&K (Shanghai, China) and prepared using acetone as a solvent. A sublethal concentration (10 ppb) of thiamethoxam (SCTH) was prepared in a 50% sucrose-water solution (1:1 w:v) with a 0.03% final concentration of acetone. A 50% sucrose-water solution (1:1 w:v) with the same concentration of acetone was also prepared as a control check (CK).

Four-day-old bees were used for the bioassays and each treatment (SCTH or CK) involved three cages of 60 bees each, all from the same colony. After ten days, we collected all bees and stored them at −80°C.

### 4.3. RNA extraction

Pools of 15 bees from each sample were prepared for total RNA extraction using Trizol reagent (Invitrogen, Carlsbad, CA, USA). RNA concentration was measured using a Qubit^®^ RNA Assay Kit in a Qubit^®^ 2.0 Fluorometer (Life Technologies, CA, USA) and RNA integrity was assessed using the RNA Nano 6000 Assay Kit of the Bioanalyzer 2100 system (Agilent Technologies, CA, USA).

### 4.4. Library preparation for transcriptome sequencing

A total of 3 *μ*g RNA was to prepare samples. Sequencing libraries were generated using a NEBNext^®^ Ultra^™^ RNA Library Prep Kit for Illumina^®^ (NEB, USA) following the manufacturer’s recommendations, and index codes were added to attribute sequences to each sample. Briefly, mRNA was purified from total RNA using poly-T oligo-attached magnetic beads. Fragmentation was carried out using divalent cations under a high temperature in NEBNext First Strand Synthesis Reaction Buffer (5 ×). First strand cDNA was synthesized using a random hexamer primer and M-MuLV Reverse Transcriptase (RNase H). Second strand cDNA synthesis was subsequently performed using DNA Polymerase I and RNase H. Remaining overhangs were converted into blunt ends via exonuclease/polymerase. After adenylation of the 3’ ends of DNA fragments, NEBNext Adaptors with hairpin loop structures were ligated to prepare for hybridization. cDNA fragments of 150-200 bp in length were selected by purifying library fragments with an AMPure XP system (Beckman Coulter, Beverly, USA). Then, 3 *μ*l USER Enzyme (NEB, USA) was used with size-selected, adaptor-ligated cDNA at 37°C for 15 min followed by 5 min at 95°C. PCR was then performed using a Phusion High-Fidelity DNA polymerase, Universal PCR primers and Index (X) Primer. Finally, PCR products were purified using an AMPure XP system and library quality was assessed using an Agilent Bioanalyzer 2100 system.

### 4.5 Clustering and sequencing

Clustering of the index-coded samples were performed on a cBot Cluster Generation System using a TruSeq PE Cluster Kit v3-cBot-HS (Illumina) according to the manufacturer’s instructions. After cluster generation, the library preparations were sequenced on an Illumina Hiseq 4000 platform and 150 bp paired-end reads were generated.

### 4.6 Read processing

Raw reads in the FASTQ format were first processed through in-house Perl scripts. Clean reads were obtained by removing low-quality reads or those containing adapters or poly-N. At the same time, the Q20, Q30 and GC contents were calculated. All downstream analyses were based on the high-quality clean reads. The index of the honeybee genome (NCBI: Amel_4.5) was built using Bowtie v2.2.3, and paired-end clean reads were aligned to the honeybee genome using TopHat v2.0.12. HTSeq v0.6.1 was used to count the reads mapped to each gene, and then the FPKM (Fragments Per Kilobase of transcript sequence per Million reads) of each gene was calculated based on the length of the gene and reads count mapped to it.

### 4.7. Differential expression analysis

Differential expression analysis was performed using the DESeq R package (2.15.3). DESeq provides statistical routines to determine differential expression in digital gene expression (DGE) data using a model based on negative binomial distribution. The resulting *P*-values were adjusted using the Benjamini and Hochberg approach for controlling the false discovery rate. Genes with an adjusted *P*-value ≤ 0.05 were considered differentially expressed. Finally, Gene Ontology (GO) enrichment analysis of differentially-expressed genes was carried out using the GOseq R package, in which gene length bias was corrected. GO terms with corrected *P*-values ≤ 0.05 were considered significantly enriched by differentially-expressed genes. In the Kyoto Encyclopedia of Genes and Genomes (KEGG) database, we used KOBAS software to test the statistical enrichment of differential gene expression in KEGG pathways.

## Author contributions

L.S.Y. and T.F.S. conceived and designed the study. T.F.S., Y.F.W and L.Q. performed the experiments. T.F.S. analyzed data and wrote the manuscript. F.L. edited and revised the manuscript. All authors have read and approved the final manuscript.

